# SOCS domain targets ECM assembly in lung fibroblasts and experimental lung fibrosis

**DOI:** 10.1101/2024.02.14.580347

**Authors:** Carina Magdaleno, Daniel J. Tschumperlin, Narendiran Rajasekaran, Archana Varadaraj

## Abstract

Idiopathic pulmonary fibrosis (IPF) is a fatal disease defined by a progressive decline in lung function due to scarring and accumulation of extracellular matrix (ECM) proteins. The SOCS (Suppressor Of Cytokine Signaling) domain is a 40 amino acid conserved domain known to form a functional ubiquitin ligase complex targeting the Von Hippel Lindau (VHL) protein for proteasomal degradation. Here we show that the SOCS conserved domain operates as a molecular tool, to disrupt collagen and fibronectin fibrils in the ECM associated with fibrotic lung myofibroblasts. Our results demonstrate that fibroblasts differentiated using TGFß, followed by transduction with the SOCS domain, exhibit significantly reduced levels of the contractile myofibroblast-marker, α-SMA. Furthermore, in support of its role to retard differentiation, we find that lung fibroblasts expressing the SOCS domain present with significantly reduced levels of α-SMA and fibrillar fibronectin after differentiation with TGFß. We show that adenoviral delivery of the SOCS domain in the fibrotic phase of experimental lung fibrosis in mice, significantly reduces collagen accumulation in disease lungs. These data underscore a novel function for the SOCS domain and its potential in ameliorating pathologic matrix deposition in lung fibroblasts and experimental lung fibrosis.

## Introduction

Idiopathic pulmonary fibrosis (IPF) is a rapidly progressive and irreversible disease with a median survival of 2.5-3.5 years (Flaherty, Thwaite et al. 2003, Perez, Rogers et al. 2003, Monaghan, Wells et al. 2004). IPF is the most common of interstitial lung disease overall, with patients experiencing progressive breathlessness, eventual respiratory failure and death. Recurrent injury to lung tissue results in a failure to resolve the wound-healing response (B, Lawson et al. 2013). The unresolved wound ultimately exhibits excessive deposition of extracellular matrix (ECM) and fibrosis leading to loss of lung function. With 34,000 new cases diagnosed each year, IPF poses a public health problem in the United States (Raghu, Weycker et al. 2006).

During fibrosis, the activation of fibroblasts to myofibroblasts, is accompanied by secretion and assembly of the ECM protein fibronectin (FN) (Muro, Moretti et al. 2008, Altrock, Sens et al. 2015). The presence of profibrotic cytokines such as TGFß in the disease milieu, further increases FN assembly/fibrillogenesis to increase ECM stiffness and cell contractility (Biernacka, Dobaczewski et al. 2011). The increased stiffness of the ECM translates to fibrogenic stimuli, in part by activation of profibrotic cytokines such as TGFß from its latent form. The active TGFß further increases matrix assembly (Wipff, Rifkin et al. 2007). The mechanical properties associated with matrix assembly modulate proliferation and differentiation of resident cells such as fibroblasts and myofibroblasts. As the accumulation of myofibroblasts and fibrosis proceeds, the tissue becomes increasingly stiff and in a pathologic feedback loop, stimulates greater ECM deposition, compromising tissue function. (Huang, Yang et al. 2012, Zhou, Huang et al. 2013, Shi, Long et al. 2014, Scott, Mair et al. 2015, Burgess, Mauad et al. 2016).

FN is a core component of the myofibroblast ECM in IPF lung tissue (Singh, Carraher et al. 2010, Tsukui, Sun et al. 2020). Like FN, fibrillar collagens are also critical and abundant ECM proteins in the fibrotic lung (Tsukui, Sun et al. 2020, Staab-Weijnitz 2022). Studies have shown that the polymerization of FN is a necessary step and a requirement that precedes collagen assembly and deposition (McDonald, Kelley et al. 1982, Velling, Risteli et al. 2002, To and Midwood 2011, Kubow, Vukmirovic et al. 2015). Incidentally, the pathological secretion and assembly of fibronectin and collagen accompanies the differentiation of fibroblasts to myofibroblasts (Muro, Moretti et al. 2008, Altrock, Sens et al. 2015). Critical to the process of FN fibrillogenesis is the Von Hippel Lindau (VHL) protein that directly binds FN to initiate polymerization of the FN protein by a process called fibrillogenesis (Ohh, Yauch et al. 1998). VHL is upregulated in IPF patients and can independently contribute to the fibrotic state when overexpressed (Zhou, Pardo et al. 2011).

Experimental lung fibrosis progresses in two discrete phases, an initial inflammatory phase with elevated levels of pro-inflammatory cytokines followed by the fibrotic phase characterized by increased expression of ECM proteins collagen and FN. Targeting the inflammatory phase using anti-inflammatory inhibitors is effective in experimental lung fibrosis, but these approaches have generally not been translatable to human disease (Raghu, Brown et al. 2008, Linke, Goren et al. 2010, Raghu, Martinez et al. 2015). Adenoviral SOCS1 (Suppressor Of Cytokine Signaling) gene transfer to the lung, 48 h prior to fibrosis disease induction (Nakashima, Yokoyama et al. 2008), ameliorates disease severity in a bleomycin-induced IPF mouse model. In this study the anti-inflammatory function of SOCS1 is exploited, prior to disease initiation, to retard the onset/progression of the disease. Whether the SOCS1 protein can also target the fibrotic phase of the disease and serve as a therapeutic intervention modality remains untested.

Within the SOCS family of proteins including SOCS 1-7 and cytokine inducible SH2-containing protein (CISH), the SOCS domain is a 40 amino acid conserved region that does not exhibit anti-inflammatory or immunosuppressive function (Piganis, De Weerd et al. 2011). We previously showed that the SOCS domain forms a Cullin-Ring ligase (CRL) complex targeting the Von-Hippel Lindau (VHL) tumor suppressor protein for ubiquitin-mediated degradation (Pozzebon, Varadaraj et al. 2013).

Here, we exploit the function of the SOCS domain in VHL degradation and demonstrate its capacity to reduce matrix assembly in myofibroblasts. Our results demonstrate that the SOCS domain significantly abrogates FN and collagen matrix assembly in differentiated lung fibroblasts and reduces the myofibroblast marker protein α-SMA. Also, we show that fibroblasts expressing the SOCS domain when differentiated using TGFß, exhibit significantly reduced myofibroblast marker proteins. In a bleomycin mouse model of pulmonary fibrosis we demonstrate that SOCS domain transduction localizes to the lung and critically, reduces collagen content in the lung. These results identify a new molecular tool and reveal new findings on the therapeutic function of the SOCS domain in ameliorating pathologic ECM deposition in the lungs.

## Materials & Methods

### Cell lines and Culture conditions

Human lung fibroblasts, IMR90, were obtained from the American Type Culture Collection (ATCC, #CCL-186) and cultured in EMEM media (ATCC, #30-2003) supplemented with 10% FBS (HyClone, #SH30109.03). Cells were maintained in a 37°C humidified incubator buffered with 5% CO_2_. All experiments were performed at 70-80% cell densities.

### Fibroblast differentiation conditions

IMR90 fibroblasts were differentiated to myofibroblasts with 5ng/mL Recombinant Human Transforming Growth Factor -β1 (TGF-β1) (Invitrogen, #PHG9204) for 48h in serum free media. TGF-β1 was reconstituted and stored according to manufacturer’s recommendations.

### Antibodies and reagents

Antibodies for immunoblotting and immunocytochemistry: GFP (Abcam, #ab183734), VHL (Cell Signaling, #68547S), HIF1-LJ (Cell Signaling Technology, #14179S), β Actin (Invitrogen, #PA5-59497), β Tubulin (Invitrogen, #MA5-16308), Fibronectin (Santa Cruz, #sc-59826), Collagen (Abcam, #ab34710), Smooth Muscle Actin (Abcam, #ab7817), Donkey Anti-Mouse Alexa Flour 568 (Invitrogen, #A10037), Donkey Anti-Rabbit Alexa Flour 568 (Invitrogen #A10042) VH298 (Axon MedChem, #2810) was used at working concentrations of 100µM for 1h in serum media.

### Adenovirus transduction

Ad-GFP (#1060), Ad-GFP #749 (amp), Ad-GFP#839 (amp) were purchased from Vector Biolabs at a viral titer of 4.5×10^10^PFU/mL, 1.2×10^11^PFU/mL, 1.1×10^12^ PFU/mL respectively. We obtained ∼80% transduction for Ad-GFP 1060 (100MOI), Ad-GFP 749 (600MOI), and Ad-GFP 839 (600MOI) at 36h.

### Protein extraction and immunoblotting

IMR90 cells were seeded on 6cm plates at a density of 163,000 cells. Protein extraction was performed at 4°C using cold SDS lysis buffer containing protease and phosphatase inhibitors (1mM DTT, 1mM EDTA, 1µg/mL Leupeptin, 100µg/mL PMSF, and 1mM Sodium Orthovanadate). Proteins were separated by SDS-Poly acrylamide gel electrophoresis and immunoblotted for specific proteins of interest. β Actin and β Tubulin were used as loading controls. Quantification of immunoblots was performed using the Li-Cor Image Studio Software version 5.2. Pixel intensities of each protein were normalized to the loading control. Three independent experiments were averaged and fold differences between fibroblasts and myofibroblasts or myofibroblasts transduced with Ad-GFP 1060 and myofibroblasts transduced with adenoviruses Ad-GFP 739 or Ad-GFP 849 were plotted and shown as bar graphs. P values were determined using Student’s t-test.

### Immunocytochemistry and imaging

For immunocytochemistry of FN, SMA, or COL, IMR90 cells were seeded on sterile coverslips in a 6-well plate at a density of 80,000 cells per well. The coverslips were fixed with 4% paraformaldehyde and permeabilized in 0.1% Triton X-100 for 1 min on ice. The cells were then blocked with 5% BSA-1x PBS for 1 h on ice. After blocking, FN antibody (1:70), SMA (1:1000) or COL (1:100) was added for 1h followed by 1h incubation with a secondary antibody. After secondary incubation, the cells were washed repeatedly with 0.2% Tween in 1xPBS. After washes, the cells were stained with the DNA stain DAPI (4, 6 diamidino-2-phenylindole dihydrochloride) (Roche, #1023627001) and mounded with Prolong gold anti-fade mound media (Invitrogen, #S36936) on glass slides. Images and z-stacks (1µM z-slice) were acquired using a Leica TCS SPEII confocal microscope at consistent acquisition parameters for each experiment.

### Fibril count and quantification

Graphs in Fig. 2b, Fig. 2c, Fig. 3b, Fig. 4b were constructed by random fields of view within each sample being used to quantify the percentage of fibril containing cells. A FN or COL track of ∼3µM in length was considered a fibril. A total of at least 25 cells were counted for each condition. Data in figures that include fibril counts are averages of 3 independent experiments. Student’s t-test was used to calculate P values.

### SMA intensities

Graphs in Fig. 2d, Fig. 3c, Fig. 4c were constructed with the intensity plugin available in Image J. Cell containing random fields of view within each sample were used to quantify the SMA intensity values. Intensities shown in figures are averages of 3 independent experiments. Student’s t-test was used to calculate P values.

### FUD and III-11C peptide treatment

FN fibril inhibitor experiments were performed by treating myofibroblasts with 500nM FUD (GenScript, #SC6004PF) or 500nM III-11C (GenScript, #SC6004PF) for 24h in serum media in addition to 5ng/mL TGF-β1. After peptide treatment, cells were fixed and processed for immunocytochemistry.

### VHL Inhibitor treatment

VHL inhibitor experiments were performed by treating myofibroblasts with 100µM VH298 for 1h in serum media. Cells were then fixed and processed for immunocytochemistry or lysed in SDS lysis buffer for immunoblotting.

### Statistical analysis

Immunoblots and immunocytochemistry experiments were performed as 3 independent trials and represented as averages ±SEM. Statistical significance was determined between fibroblasts and myofibroblasts or myofibroblasts transduced with Ad-GFP 1060 and myofibroblasts transduced with Ad-GFP 749 and Ad-GFP 839 for FN/COL. Statistical significance for fibrils counts, SMA intensities and immunoblot intensities were calculated using Student’s t-test. Statistical significance (P-value) was determined using GraphPad Prism 8.

### Animal Models

All protocols concerning animal use were approved by the Institutional Animal Care and Use Committees at the Northern Arizona University and conducted in strict accordance with the National Institutes of Health Guide for the Care and Use of Laboratory Animals. Studies were conducted with 10-12 week old C57Bl/6 mice. Mice were housed in a temperature and humidity-controlled pathogen-free facility with a 12:12 hour light:dark cycle. Animals were housed within a limited access rodent facility and kept in groups of a maximum of 4 mice per cage. Mice were housed in polypropylene cages with solid bottoms and wood shavings or corn cobb as bedding material. At study termination, mice were euthanized via anesthesia overdose and exsanguination.

### Bleomycin lung fibrosis model and viral transduction

IPF was established in C57Bl/6 mice (10-12 weeks) by oral aspiration (OA) of bleomycin (3.07 units/kg) on Day 0. Adenovirus (Ad-GFP-Empty, SOCS domain and SOCS domain mutant) was administered intranasally (IN) on Day 9. Animals were euthanized on Day 21 (5 animals in each group).

Collagen content (hydroxyproline assay) in the lung was determined from lung lysates collected on Day 21 of the study.

### Immunohistochemistry and GFP-DAB staining and quantification

To determine optimum adenoviral titer for transduction, C57BL/6 mice were administered adenovirus 10^8^ PFU and 10^9^PFU or vehicle control (saline) on Day 0. On Day 3 following euthanasia, the lungs were inflated and fixed in formalin 10%. Tissues were processed in paraffin, sectioned and stained using GFP-DAB for assessment of transduction and H&E. Images were imported in HALO 3.0, regions of interest (lung tissue) annotated, and a quantitative image analysis algorithm performed. The algorithm (Cytonuclear V1.6) was used to segment nuclei based on hematoxylin staining, classify cells as either positive or negative for DAB staining and stratify positively stained cells as being 1+, 2+ or 3+ staining intensity (1+ being weak, 2+ being moderate, 3+ being strong staining intensity). To assign the H-score, a classification system was used to assess the total number of positive cells as well as the intensity of staining of each positive cell (on a scale of 0-3). The formula is calculated as [0*(ratio of negative cells)+1*(ratio of 1+ cells)+2*(ratio of 2+ cells)+3*(ratio of 3+ cells)].

## Results

### SOCS domain decreases VHL levels in lung myofibroblasts

We previously showed that a 40 amino acid conserved domain in the SOCS family of proteins called the SOCS domain, degrades VHL, unlike a SOCS domain mutant that is deficient in interacting with the CRL complex (Pozzebon, Varadaraj et al. 2013). To confirm VHL degradation in myofibroblasts, in the presence of the SOCS domain, we differentiated IMR90 lung fibroblasts with TGFβ for 48 h and transduced cells with adenovirus expressing the SOCS domain, SOCS domain mutant or empty vector. Viral transduction of differentiated myofibroblast cells was confirmed by detectable GFP reporter expression (**Fig 1A**). Specifically, in cells transduced with the SOCS domain, VHL depletion was observed (**Fig 1B**, **Fig 1C**). Since it is well established that the depletion of VHL preserves HIF-1α protein stability (Mack, Rathmell et al. 2003), we therefore examined HIF-1α levels under the above-mentioned conditions. We detected significantly increased HIF-1α protein levels that corresponded to depleted VHL protein, in the SOCS domain transduced cells (**Fig 1D**, **Fig 1E**). There was no significant difference in HIF-1α protein levels between differentiated myofibroblasts versus cells transduced with the SOCS domain mutant or empty virus. Together, the results demonstrate that the SOCS domain (unlike the mutant) significantly reduces VHL protein levels in differentiated lung fibroblast cells.

**Fig 1:**
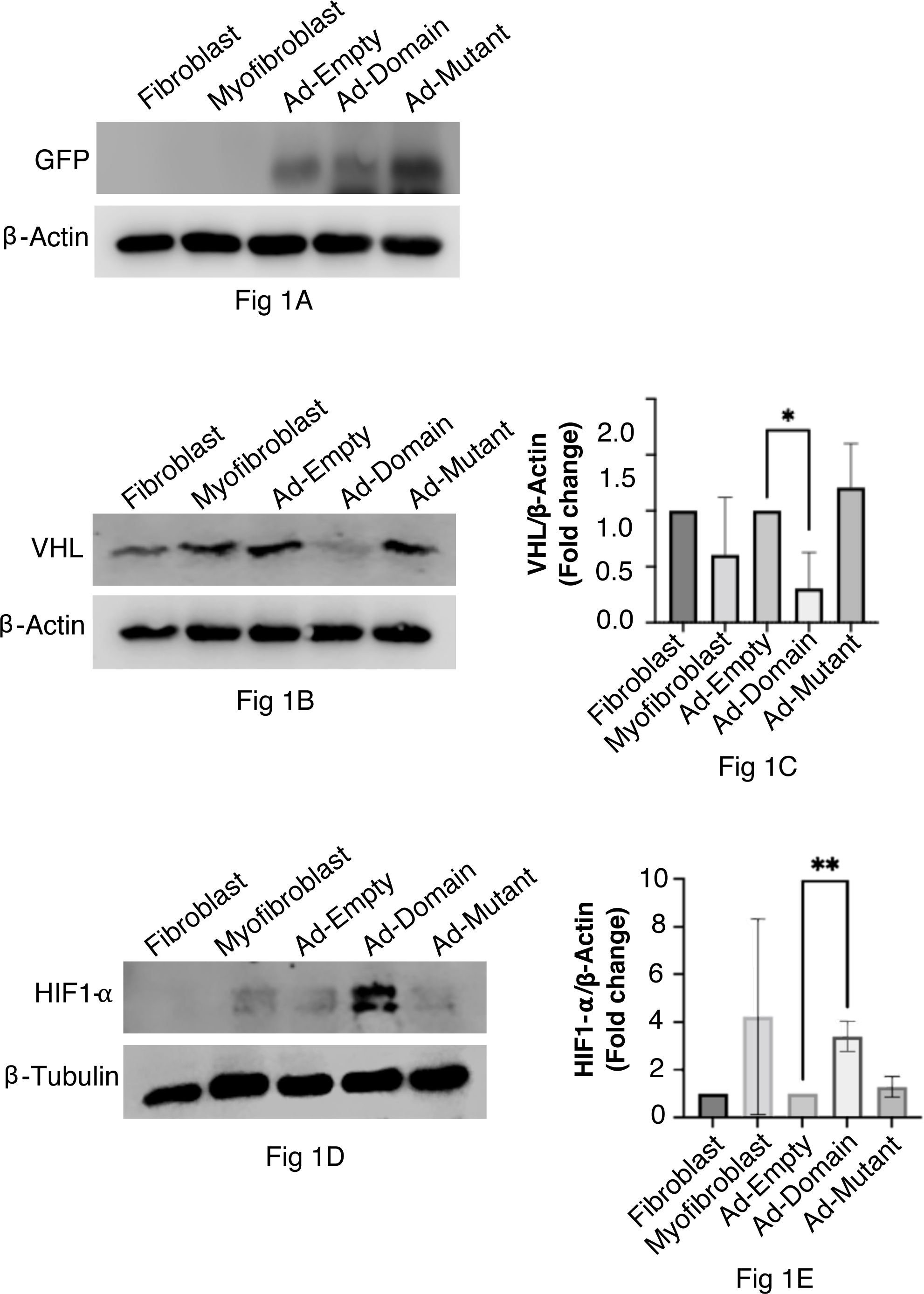
SOCS domain degrades VHL protein in myofibroblasts. (A,B,D) IMR90 lung fibroblasts were untreated or differentiated with 5ng/mL TGFβ for 48 h. Differentiated myofibroblasts were then transduced with Ad-Empty 100 MOI, Ad-Domain 600 MOI or Ad-Mutant 600 MOI for ∼36 h. Cells were lysed in SDS lysis buffer and immunoblotted for GFP, VHL and HIF1-LJ. β-Actin and β-Tubulin were used as the loading controls. (C&E) Quantification of VHL and HIF1-LJ normalized to their respective loading controls, on immunoblots are represented as averages of three independent trials. Statistical significance between fibroblasts and myofibroblasts and Ad-Empty and Ad-Domain or Ad-Mutant P-values were determined using Student’s t-test.

### SOCS domain reduces matrix assembly and decreases αSMA in myofibroblasts

Prior studies have established the contribution of excess deposition of matrix proteins collagen and FN in preserving a fibrotic state (Kuhn, Boldt et al. 1989, Herrera, Henke et al. 2018). FN is a core matrisome protein, requiring VHL for its matrix formation (Ohh, Yauch et al. 1998). Therefore, we hypothesized that VHL degradation using the SOCS domain might abrogate matrix formation. To test this (**Fig 2A**), we differentiated fibroblasts using TGFβ. TGFβ treatment increased FN and collagen matrix formation and increased levels of α-smooth muscle actin (SMA), a marker for contractile myofibroblasts. Transduction of myofibroblasts with the SOCS domain resulted in a significant decrease in the assembly of ECM proteins FN and collagen, as observed by the decrease in the length of the matrix fibrils (**Fig 2B**, **Fig 2C**). Interestingly, in these conditions, we also detected a significant decrease in levels of α-SMA (**Fig 2D**). In line with these data, we observed no significant decrease in fibril assembly of matrix proteins or α-SMA levels in myofibroblasts transduced with the SOCS domain mutant. Quantification of the transduced cells (as determined from GFP reporter expression) confirmed the significant decrease in matrix fibrils fibronectin and collagen, as well as levels of α-SMA, in cells expressing the SOCS domain (**Fig 2B-D**).

**Fig. 2:**
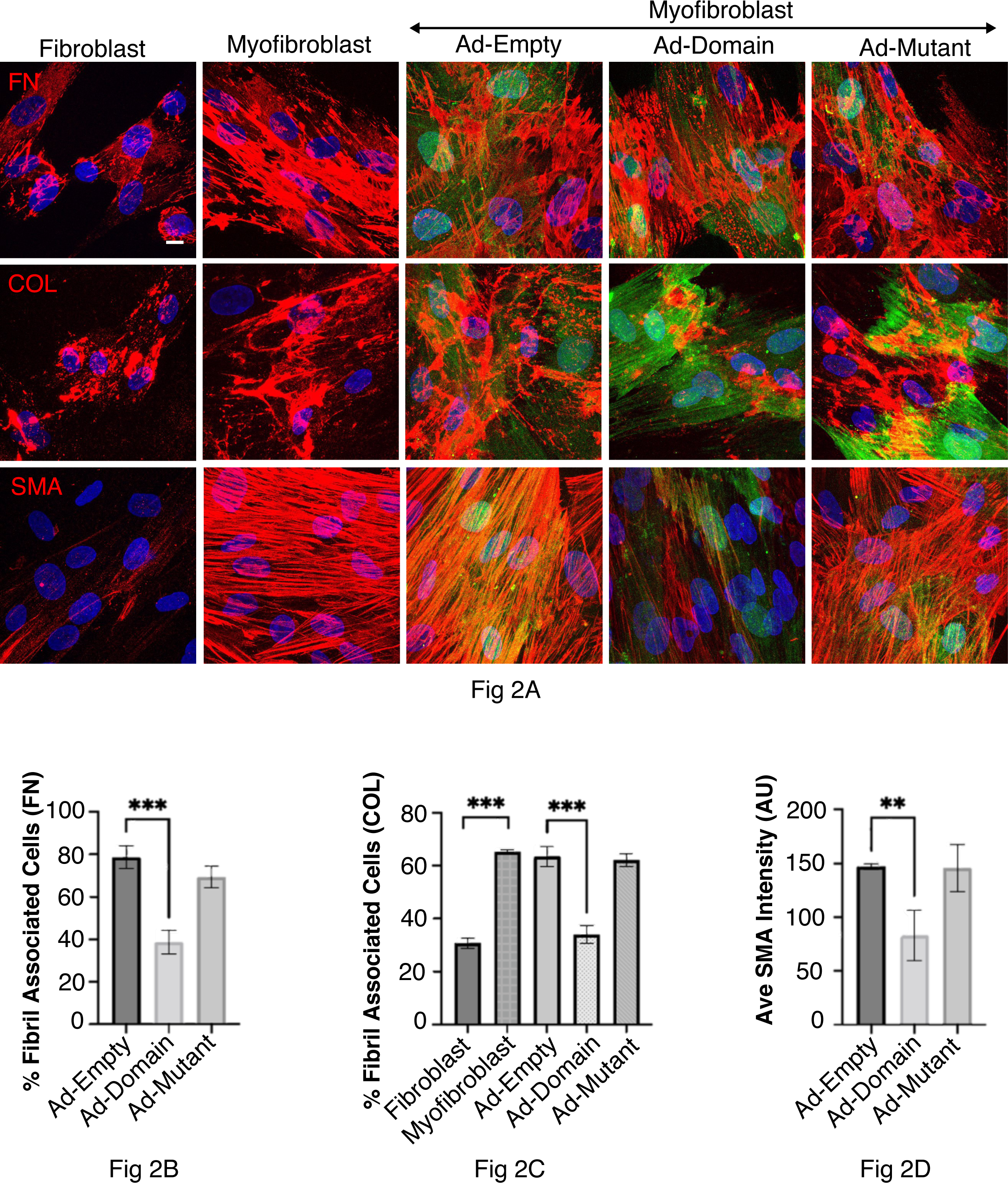
SOCS domain fragments FN matrix and reduces levels of SMA protein in myofibroblasts. (A) IMR90 lung fibroblasts were untreated or differentiated with 5ng/mL TGFβ for 48 h. Differentiated myofibroblasts were then transduced with Ad-Empty (100 MOI), Ad-Domain (600 MOI) or Ad-Mutant (600 MOI) for ∼36 h and immunostained for FN (red), COL (red) or SMA (red) and nuclear stain DAPI (blue). Myofibroblasts transduced with Ad-Empty (100 MOI), Ad-Domain (600 MOI) or Ad-Mutant (600 MOI) appear green (GFP reporter). Scale bar =10µM. (B&C) GFP reporter-expressing transduced cells as in (A) were quantified for percent of fibronectin fibril-containing cells and collagen fibril-containing cells respectively. Bar graphs are averages of three independent experiments. Student t-test was used to calculate P values. (D) Transduced cells expressing the GFP reporter as in (A) were used to quantify the SMA intensity values. Intensities shown in the graph are averages of three independent experiments. Student’s t-test was used to calculate P values.

### SOCS domain retards fibroblast differentiation

Having confirmed that the SOCS domain lowers the levels of myofibroblast marker proteins, despite differentiation by TGFβ, we next evaluated whether the SOCS domain protects against differentiation by TGFβ. For this we transduced fibroblast cells with adenovirus expressing the SOCS domain, mutant or empty vector, prior to differentiation with TGFβ (**Fig 3**). Untransduced cells when differentiated by TGFβ treatment, exhibited a noticeable increase in FN matrix deposition and α-SMA levels. However, fibroblast cells transduced with the SOCS domain failed to upregulate FN matrix formation or α-SMA levels when differentiated with TGFβ (**Fig 3A**). Quantification of transduced cells containing a fibronectin matrix, revealed a significant decrease in the numbers of these cells in the presence of the SOCS domain (**Fig 3B**). In line with this data, α-SMA levels in the SOCS domain expressing cells significantly decreased compared to cells expressing either the empty virus or the mutant domain (**Fig 3C**). These data demonstrate that SOCS domain expression retards fibroblast differentiation.

**Fig. 3:**
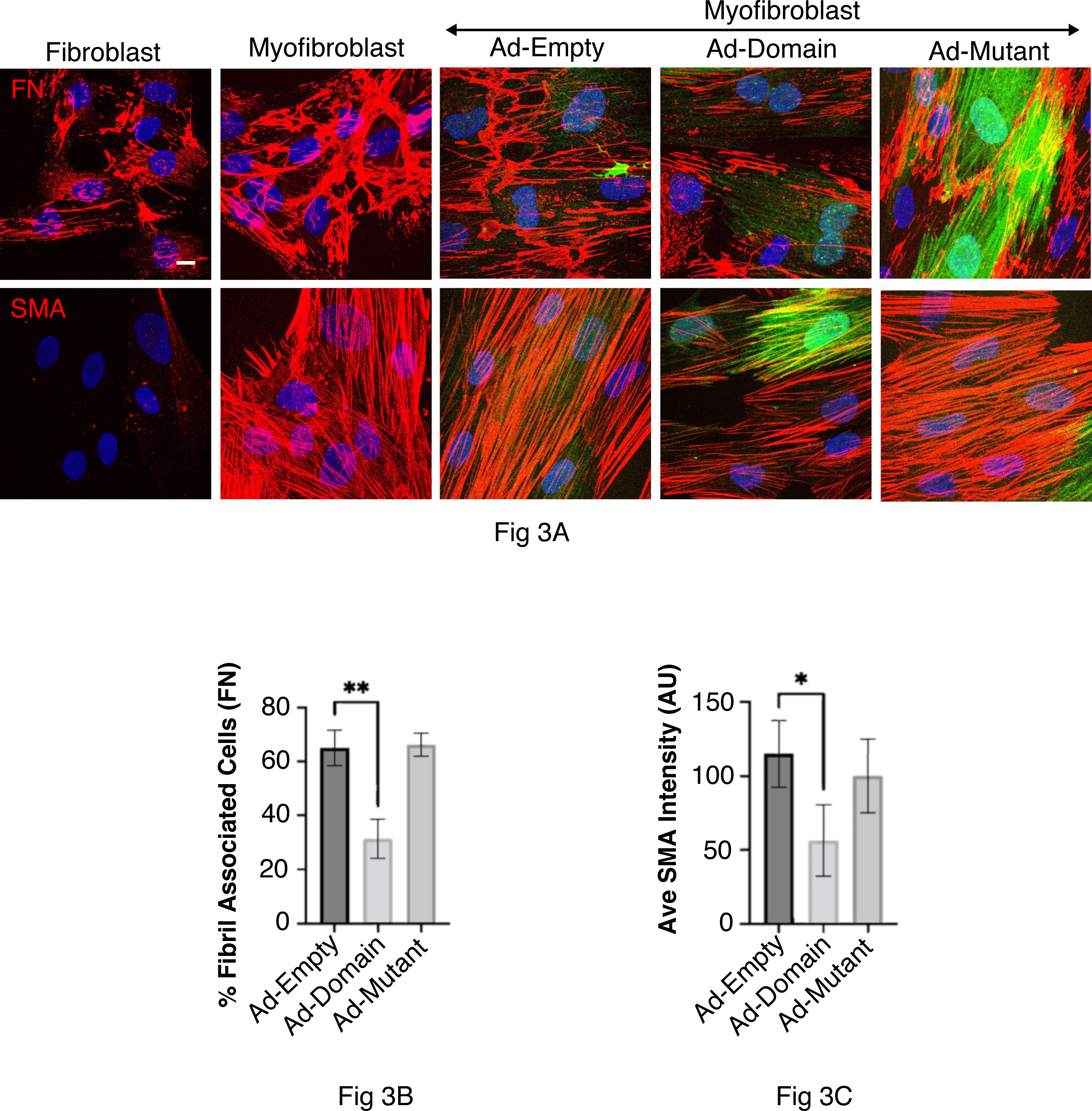
SOCS domain reduces differentiation of fibroblasts to myofibroblasts. (A) IMR90 lung fibroblasts were untreated or transduced with Ad-Empty (100 MOI), Ad-Domain (600 MOI) or Ad-Mutant (600 MOI) for ∼36 h. Cells were then differentiated with 5ng/mL TGFβ for 48 h. Fibroblasts, myofibroblasts (untransduced) and transduced myofibroblasts were immunostained for FN (red) or SMA (red) and nuclear stain DAPI (blue). Myofibroblasts transduced with Ad-Empty (100 MOI), Ad-Domain (600 MOI) or Ad-Mutant (600 MOI) appear green (GFP reporter). Scale bar =10µM. (B) Transduced cells as in (A) were quantified for percent of fibronectin fibril-containing cells and represented as bar graphs. Only cells that expressed the GFP reporter were included in the analysis. (C) Transduced cells expressing the GFP reporter were used to quantify the SMA intensity values. Intensities shown in the graph are averages of three independent experiments. Student’s t-test was used to calculate P values.

### Matrix disassembly is insufficient in reversing myofibroblast marker proteins

The function of the SOCS domain in decreasing matrix deposition of myofibroblasts, led us to test whether matrix disruption was sufficient to reverse the myofibroblast marker α-SMA. To test this, we blocked fibronectin fibril assembly using a 49mer peptide (functional upstream domain, FUD) derived from the *Streptococcus pyogenes* adhesion F1 protein, that binds to the N-terminal domain of FN, the region which is responsible for fibril formation. Since FUD does not diminish FN protein levels, this approach specifically targets fibril assembly without altering the protein composition of the ECM (McKeown-Longo and Mosher 1985, Chernousov, Fogerty et al. 1991, Sechler, Takada et al. 1996, Tomasini-Johansson, Kaufman et al. 2001, Tomasini-Johansson, Annis et al. 2006). As was expected, treatment of myofibroblasts with the FUD peptide abrogated fibril assembly (unlike treatment with the control peptide III-11C) (**Fig 4A&B**). Despite this, we observed no concomitant change in levels of α-SMA in these cells (**Fig 4A&C**). These data suggest that matrix disruption alone is insufficient to reduce myofibroblast marker proteins.

**Fig. 4:**
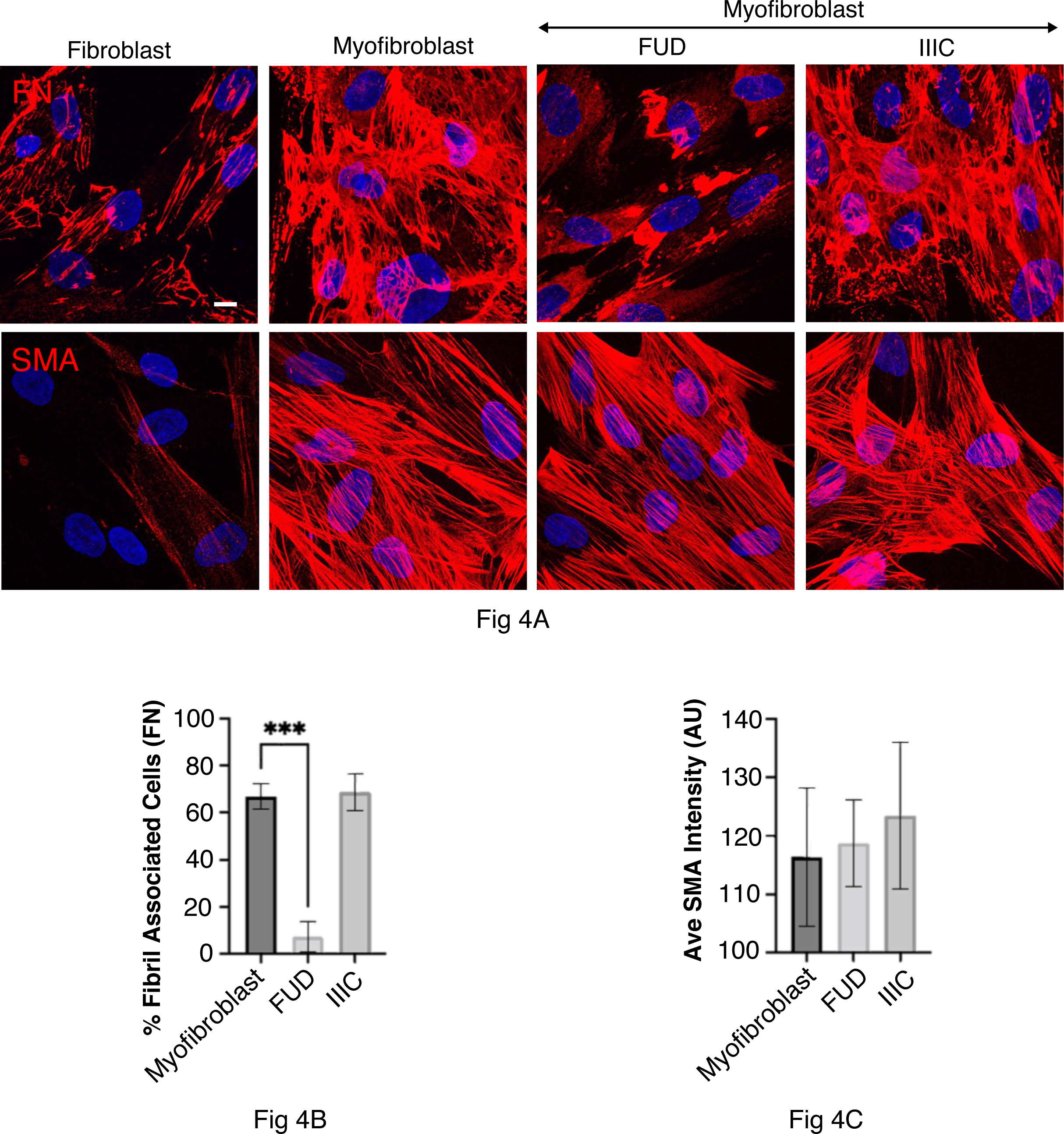
FN fibril inhibition does not decrease SMA intensity in myofibroblasts. (A) IMR90 lung fibroblasts were untreated or differentiated with 5ng/mL TGFβ. Differentiated myofibroblasts were then treated with 500nM FUD or 500nM IIIC-11C for 24 h in addition to 5ng/mL TGFβ. Cells were immunostained for FN (red) or SMA (red) and nuclear stain, DAPI (blue). Scale bar =10µM. (B) Myofibroblasts and myofibroblasts treated with FUD or IIIC were quantified for fibronectin fibril-containing cells or (C) used to quantify the SMA intensity values. Bar graphs are averages of three independent trials. Student’s t-test was used to determine P values.

### VHL inhibition does not reduce myofibroblast marker proteins

We previously showed that the SOCS domain forms an E3 ubiquitin ligase complex and degrades VHL (Pozzebon, Varadaraj et al. 2013). VHL degradation in myofibroblast cells coincides with HIF1α stabilization. To specifically investigate whether HIF1α stabilization by the SOCS domain might be a possible mechanism for reversing markers of myofibroblasts, we first inhibited VHL using VH298 a chemical probe reported to inhibit VHL by preventing the VHL-HIF1α interaction (Frost, Galdeano et al. 2016). Although treatment of myofibroblasts with the VH298 inhibitor increased HIF1α levels, as was expected (**Fig 5A**), we observed no change in FN matrix assembly and levels of α-SMA in these cells (**Fig 5B**). This data suggests that inhibition of VHL and HIF1α stabilization alone is insufficient in reducing the α-SMA levels in myofibroblast cells.

**Fig. 5:**
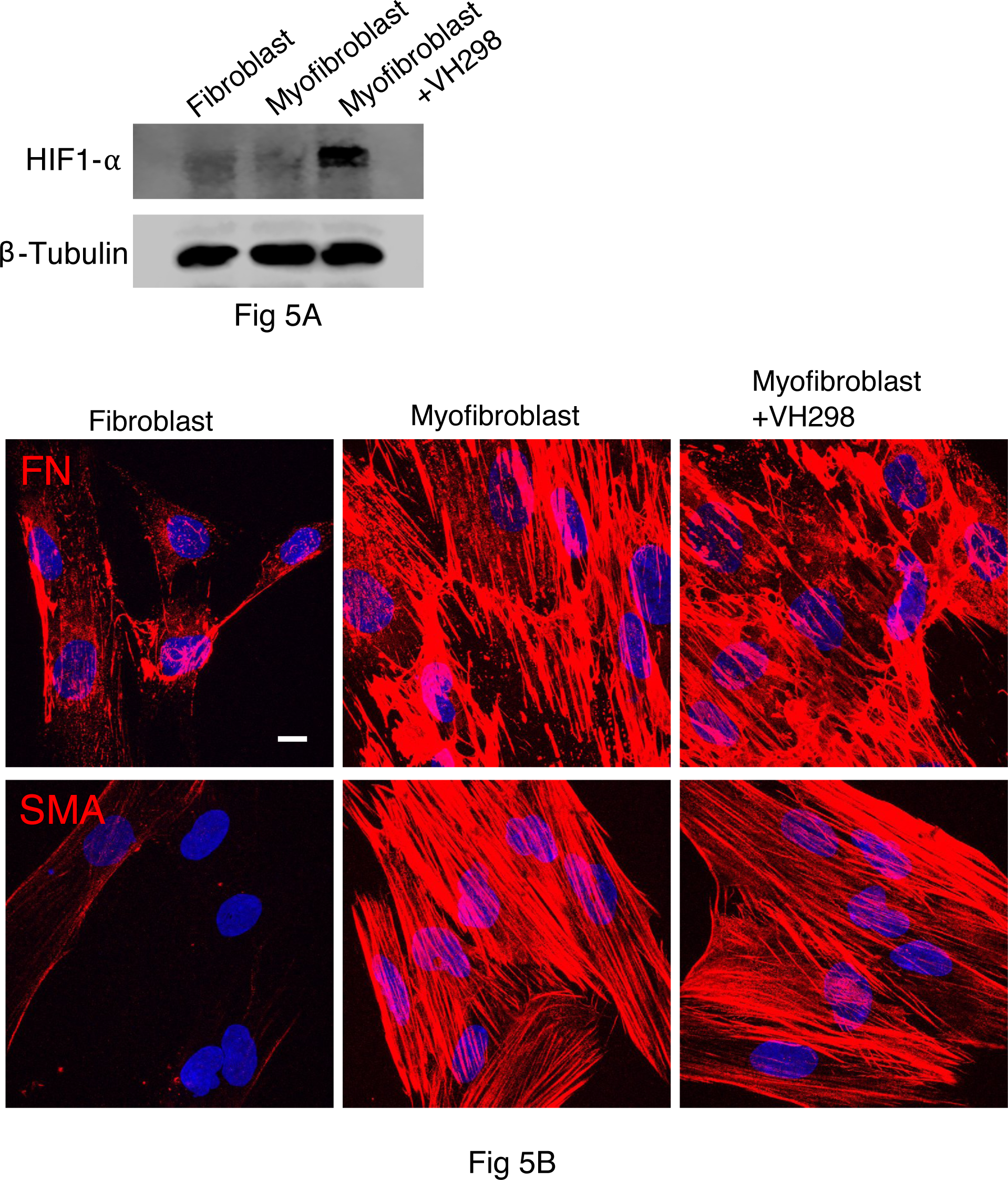
VHL protein inhibition does not decrease SMA intensity in myofibroblasts. (A) IMR90 lung fibroblasts were untreated or differentiated with 5ng/mL TGFβ for 48 h. Differentiated myofibroblasts were then treated with 100µM VH298 for 1 h. Cells were then lysed in SDS buffer and immunoblotted for HIF1-LJ. β-Tubulin was used as the loading control. (B) Fibroblasts, myofibroblasts and myofibroblasts treated as in (A) were immunostained for FN (red) or SMA (red) and nuclear stain, DAPI (blue). Scale bar =10µM.

### SOCS domain significantly reduces collagen content in in vivo model of bleomycin-induced IPF

Our in vitro findings demonstrated the reduction of ECM and myofibroblast marker proteins by the SOCS domain. To investigate the protective function of the domain in vivo, we first optimized the localization and expression of adenovirus vectors to mouse lungs in vivo. Adenovirus-mediated transduction to mouse lungs was confirmed by intranasal delivery of empty adenovirus vector expressing the GFP reporter. As shown in **Fig 6A-C**, viral titers of 1×10^8^ PFU and 1×10^9^ PFU were delivered on Day 0 and mouse lung tissue was evaluated on Day 5 for GFP-DAB levels by immunohistochemistry. Substantial GFP expression was detected at 1×10^9^ PFU compared to the negligible staining at 1×10^8^ PFU. GFP expression was restricted to the lung tissue and we observed no expression in the diaphragmatic smooth muscle in close proximity to the lungs and no expression was noted in the lymphoid tissue found within and near the lungs. Quantification of the GFP intensities (H-score) (**Fig 6C**) confirmed significant increase in GFP levels at the higher viral titer of 1×10^9^ PFU. This data confirmed localization and optimum viral titer for transduction to the mouse lung.

**Fig. 6:**
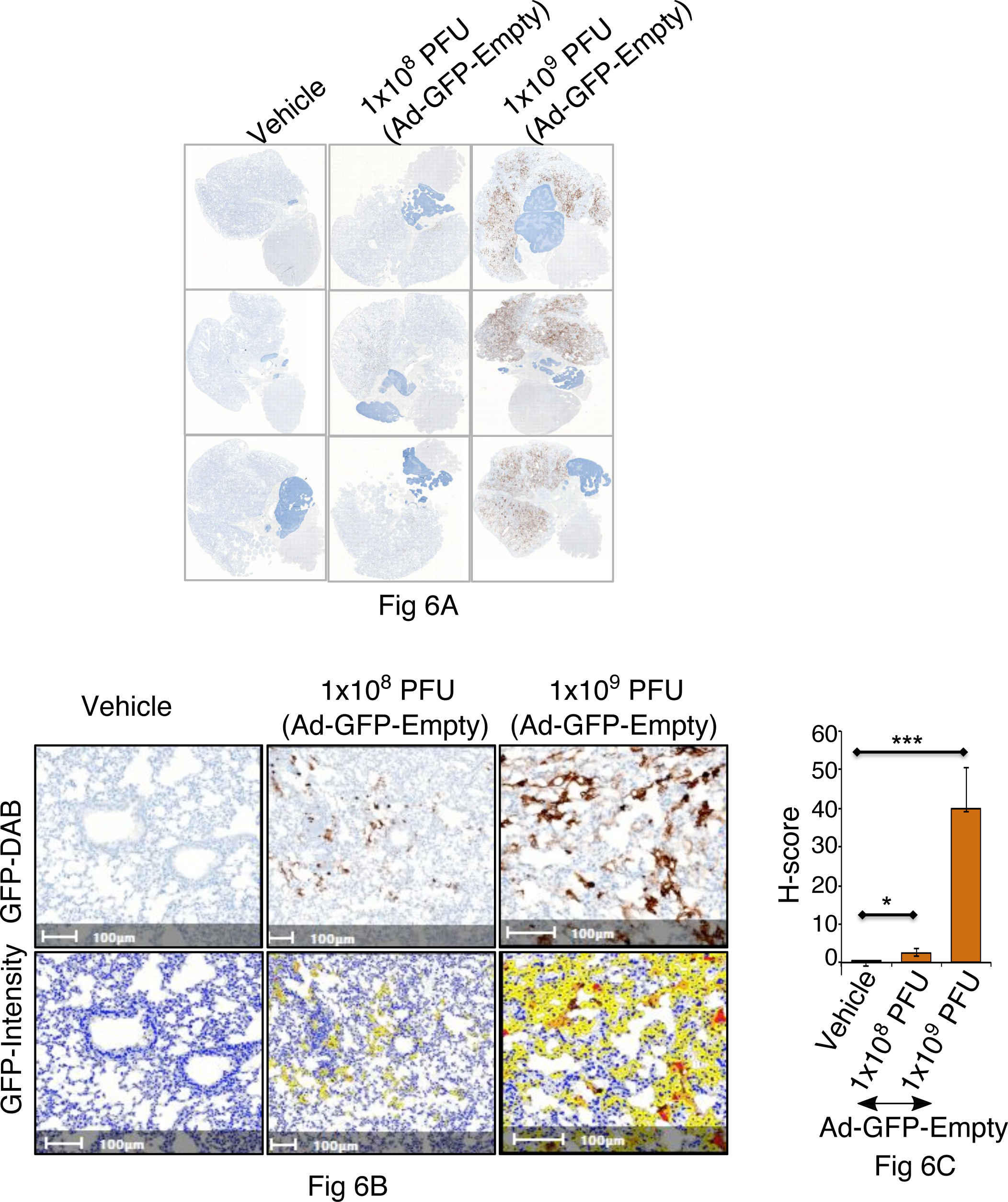
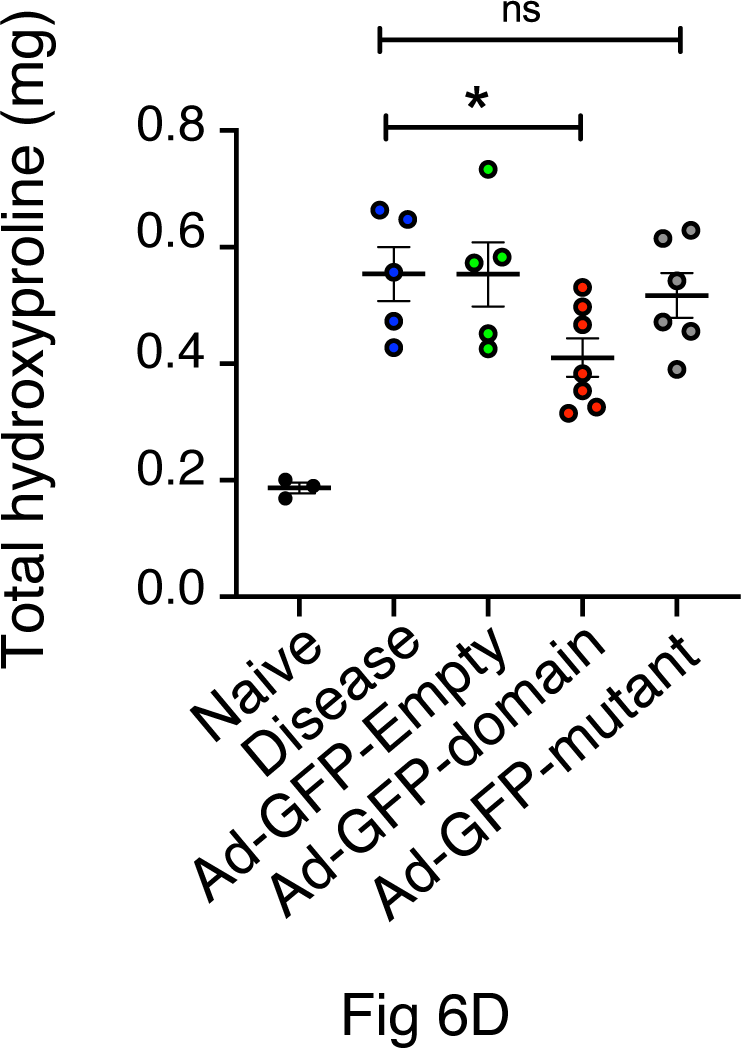
Intranasal delivery of Adenoviral-GFP is localized to lung tissue and SOCS domain delivery significantly reduces collagen deposition in fibrotic lung. (A) C57BL/6 mice were intranasally administered vehicle (phosphate buffered saline) and adenoviral-GFP on Day 0 at 10^8^ PFU and 10^9^ PFU as depicted. Lung tissue was collected on Day 5 and Formalin Fixed paraffin embedded (FFPE) lung tissue was subjected to immunohistochemistry (IHC) staining using GFP-DAB antibody. Increased transduction of Ad-GFP at 10^9^ PFU was observed compared to transduction at 10^8^ PFU 5 days following intranasal delivery of Ad-GFP and vehicle control (n=3). Magnification=20x. Scale bar=10μm. (B) FFPE lung tissue sections analyzed for GFP expression. Top row depicts GFP-DAB staining and bottom row shows nuclei (blue). (C) Positive stain intensity from the bottom row in FIG. 6B, assigned using the scores: 1+ yellow, 2+ orange and 3+ red was used to quantify extent and intensity of GFP represented using the histoscore (H-score). Bar graph represents mean ± SD, n=3. Statistical significance between vehicle (phosphate buffered saline) versus transduced and P-values (*P<0.05, ***P<0.001) are depicted. (D) C57BL/6 mice were administered bleomycin 3.07 units/kg by oropharyngeal aspiration on D0 and intranasally administered adenovirus on D9. Lung lobes collected on D21 was quantified for collagen deposits (hydroxyproline assay). Disease vs SOCS domain *p<0.05. n=6 mice per group.

Next, pulmonary fibrosis was established in a bleomycin (BLM) mouse model by delivery of bleomycin 3.07 units/kg on Day 0. On Day 9, during the fibrotic phase of the disease, vehicle (saline) and adenovirus (SOCS domain, SOCS domain mutant and Empty vector) was administered intranasally at a viral titer of 1×10^9^ PFU. On Day 21 after initiation of bleomycin injury, mouse lung tissue was collected, lysed and analyzed for collagen content using the biochemical hydroxyproline assay (**Fig 6D**). As was expected, collagen content was elevated in the disease lung. Lung tissue transduced with the Empty vector and SOCS domain mutant also exhibited elevated collagen content that was comparable to levels in the disease lung. However, a significant decrease in the hydroxyproline content was observed in mouse lung transduced with the SOCS domain. These data confirm the role of the SOCS domain in specifically reducing the structural collagen that is associated with IPF disease severity.

## Discussion

In this study we establish the role of the 40 amino acid SOCS domain in reducing FN and collagen matrix assembly in differentiated lung fibroblasts. Our results confirm the specific function of the SOCS domain in reducing matrix assembly in myofibroblasts, cells that would otherwise exhibit abundant levels of ECM upon differentiation in the presence of TGFß (**Fig 2A-C**). Moreover, the significantly lower levels of α-SMA in myofibroblasts expressing the SOCS domain, confirms that the domain is capable of influencing the differentiation state of the cell (**Fig 2A and 2D**). Myofibroblast dedifferentiation has been reported in response to several inhibitors targeting FN matrix assembly. For instance, the Fn52 antibody targeted against the N-terminal 30 kDa region that is required for FN self-association, reduces the fibrotic features of lens epithelial cells (Tiwari, Kumar et al. 2016). Similarly, inhibition of the Rho-ROCK pathway, required for FN assembly has been reported to attenuate the contractile markers of myofibroblasts (Zhong, Chrzanowska-Wodnicka et al. 1998, Huang, Yang et al. 2012, Marinkovic, Liu et al. 2013, Zhou, Huang et al. 2013, Tiwari, Kumar et al. 2016) and plasminogen-mediated FN matrix degradation has been shown to trigger apoptotic clearance of myofibroblasts (Horowitz, Rogers et al. 2008). Clearly, these studies emphasize matrix restructuring as drivers of the dedifferentiation program, and our results demonstrate the capacity of SOCS domain transduction to engage similar fibroblast behaviors.

While our data in **Figs 2&3** suggested that the SOCS domain reverses the myofibroblast phenotype through the degradation of the FN matrix, we found that additional pathways triggered by the SOCS domain are likely necessary for dedifferentiation. Notably, although treatment of myofibroblast cells with the FN matrix disrupting peptide FUD significantly reduced matrix assembly, we observed no concomitant change in levels of the myofibroblast marker protein α-SMA, dissociating matrix levels of myofibroblasts from its contractile phenotype associated with increased levels of α-SMA (**Fig 4**). Therefore, one of the potential mechanisms of dedifferentiation may involve VHL. Clearly, the SOCS domain, which we have shown in previous work to form a functional CRL complex to degrade VHL (Pozzebon, Varadaraj et al. 2013), in this study elicits dedifferentiation, unlike the SOCS domain mutant that is unable to degrade VHL (**Fig 1B and Fig 1C**). However, we found that VHL inhibition using VH298 was unable to restore a fibroblast-like phenotype (**Fig 5**). One possible explanation for this divergence may be due to distinct dedifferentiation functions associated with VHL degradation as opposed to VHL inhibition. Whilst VHL inhibition using VH298 prevents VHL-HIF-1α interaction and stabilizes HIF-1α (**Fig 5A**), the VH298 inhibitor does not inhibit the HIF-1α independent functions of VHL (Frost, Rocha et al. 2021). Instead, VHL degradation that is initiated by SOCS domain expression would abrogate both the HIF-1α -dependent and independent roles of VHL that is not phenocopied by treatment with the VHL inhibitor and therefore the consequence of HIF-directed effector pathways (Frost, Rocha et al. 2021). Several proteins are regulated by VHL in a HIF-independent manner by direct interaction and ubiquitination - epidermal growth factor receptor (Zhou and Yang 2011), atypical protein kinase C (Okuda, Saitoh et al. 2001), Sprouty2 (Anderson, Nordquist et al. 2011), b-adrenergic receptor II (Xie, Xiao et al. 2009), Myb-binding protein 160 (Lai, Qiao et al. 2011), RNA polymerase subunits Rpb1 (Kuznetsova, Meller et al. 2003, Mikhaylova, Ignacak et al. 2008) and Rpb7 (Na, Duan et al. 2003) and proteins related to spindle orientation and microtubule stability (Hergovich, Lisztwan et al. 2003, Thoma, Toso et al. 2009). Although this study does not provide the precise signaling events activated by the SOCS domain to initiate dedifferentiation, the data we have, point to a requirement for both matrix disassembly and VHL degradation (**Figs 2&3**).

The physiological relevance of in vitro effects of the SOCS domain are well supported by the significantly lower levels of collagen observed in lungs of bleomycin-treated mice transduced with the SOCS domain and not the SOCS domain mutant (**Fig 6**). Collagen accumulation is a critical clinicopathological feature associated with perturbation of collagen homeostasis in fibrotic lung disease. Although recent work using scRNAseq and proximity ligation in situ hybridization (PLISH) has identified several subsets of myofibroblast cells in the disease lung, all subsets of these cells are characterized by abundant collagen accumulation (Tsukui, Sun et al. 2020). Our in vitro and in vivo data demonstrating the significant decrease in collagen levels upon SOCS domain expression underscores the therapeutic potential of the peptide.

One of the limitations of our findings is the medium to low transduction efficiency of the adenoviral vector, which we predict may be 40% or lower in the mouse lung (**Fig 6A**). We acknowledge that single cell analyses of fibroblast subpopulations expressing the SOCS domain will be necessary in the future to molecularly assess dedifferentiation mechanisms in vivo. Alternatively, more efficient peptide delivery methods may be necessary to investigate these mechanisms further.

IPF has very few effective treatment options. As neither of the current therapies for IPF target matrix reorganization, our results highlight additional pathways -matrix reorganization and VHL-mediated pathways combined, that warrant further investigation in the development of novel therapeutics for IPF.

## Acknowledgements

We thank Prof. Jane Sottile (University of Rochester) for generously sharing the FUD and control peptide III-11C. Funding support was provided by the National Institutes of Health grants 1R15HL154051-01A1 to AV and NR.

## Data sharing

The data that support the findings of this study are available from the corresponding author upon reasonable request.

